# Deep evolution of MADS-box genes in Archaeplastida

**DOI:** 10.1101/2023.02.13.528266

**Authors:** Lydia Gramzow, Chiara Tessari, Florian Rümpler, Günter Theißen

## Abstract

MADS-box genes represent a paneukaryotic gene family encoding transcription factors. Given its importance for essential functions in plants, animals and fungi, such as development of organ identity and mating type determination, the phylogeny of MADS-box genes is of great biological interest. It has been well established that a gene duplication in the stem group of extant eukaryotes generated two clades of MADS-box genes, termed Type I and Type II genes. Almost all Type II genes of land plants contain a keratin-like (K) domain in addition to the family-defining, DNA-binding MADS (M) domain and are also termed MIKC-type genes. Due to a lack of sampling of MADS-box genes in Archaeplastida (rhodophytes, glaucophytes, chlorophytes, and streptophytes) except land plants, the deep evolution of MADS-box genes in plants remains poorly understood, however. Here we use the genomic and transcriptomic ressources that have become available in recent years to answer longstanding questions of MADS-box gene evolution in Archaeplastida. Our results reveal that archaeplastid algae likely do not harbour Type I MADS-box genes. However, rhodophytes, glaucophytes, prasinodermophytes and chlorophytes possess Type II MADS-box genes without a K domain. Type II MADS-box genes with a K domain are found only in streptophytes. This corroborates previous views that some Type II gene acquired a K domain in the stem group of extant streptophytes, generating MIKC-type genes. Interestingly, we found both variants of Type II genes - with (MIKC) and without a K domain - in streptophyte algae, but not in land plants (embryophytes), suggesting that Type II genes without a K domain (ancestral Type II genes) were lost in the stem group of land plants. Our data reveal that the deep evolution of MADS-box genes in “plants” (Archaeplastida) was more complex than has previously been thought.

## Introduction

### MADS-box genes

MADS-box genes encode for MADS-domain transcription factors that share the conserved DNA-binding MADS domain (Messenguy and Dubois 2003). Henceforth, we use the term “box” for DNA and “domain” for protein sequence. The MADS domain is named after four of the first five proteins identified to harbour this domain: MINICHROMOSOME MAINTENANCE 1 of yeast, AGAMOUS of thale cress, DEFICIENS of snap dragon and SERUM RESPONSE FACTOR of humans (Schwarz-Sommer, et al. 1990). Subsequently, many more MADS-box genes have been identified from a diversity of eukaryotes, but not in prokaryotes, suggesting that this gene family evolved in the stem group of extant eukaryotes (Gramzow, et al. 2010). Furthermore, two types of MADS-box genes have been classified in animals and fungi, the MEF2- and SRF-like genes (Shore and Sharrocks 1995). Similarly, Type I and Type II MADS-box genes are distinguished in plants (Alvarez-Buylla, et al. 2000). It has been hypothesized that Type I MADS-box genes are orthologous to SRF-like genes and Type II genes are orthologous to MEF2-like genes (Alvarez-Buylla, et al. 2000; Becker and Theißen 2003; Gramzow, et al. 2010). Hence, we will collectively refer to both, SRF-like and Type I genes of plants as Type I genes and to MEF2-like as well as Type II genes of plants as Type II genes. The Type II MADS-box genes of land plants encode a second conserved domain which was termed Keratin-like (K) domain due to its similarity (but not homology) to keratin (Ma, et al. 1991). The K domain follows after the MADS (M) and an intervening (I) domain and is followed by the C-terminal (C) domain. Due to this encoded domain structure, Type II MADS-box genes of plants have also been termed MIKC-type genes (Münster, et al. 1997).

MADS-box genes have been functionally characterized in amoeba, animals, fungi and plants (e.g. Elble and Tye 1992; Molkentin and Olson 1996; Arsenian, et al. 1998; Galardi-Castilla, et al. 2013). MADS-box genes of amoeba are involved in cell-differentiation during multicellular growth (Escalante, et al. 2001; Galardi-Castilla, et al. 2013). Similarly, animal MADS-box genes have been shown to be crucial for the differentiation of muscle and nerve cells (reviewed in Potthoff and Olson 2007; Knöll and Nordheim 2009). In fungi, the role of MADS-box genes may be more diverse, with functions e.g. in cell-cycle progression (Elble and Tye 1992), cell-wall integrity (Damveld, et al. 2005) and arginine metabolism (Messenguy and Dubois 1993). MADS-box genes in land plants, especially flowering plants, have been well studied and shown to be involved in almost all developmental processes, most prominently in the development of flowers (Gramzow and Theissen 2010; Smaczniak, et al. 2012). However, studies on the function and evolution of MADS-box genes in Archaeplastida other than land plants remain scarce.

### Archaeplastida

Archaeplastida is a potential clade comprised of mainly photosynthetic eukaryotes (Bowles, et al. 2022). Archaeplastida formed when an ancestral eukaryote acquired a chloroplast by the primary endosymbiosis of an ancestral cyanobacterium (Ponce-Toledo, et al. 2017). Since their origin in the paleoproterozoic era about 2 billion years ago, Archaeplastida have greatly shaped the appearance of planet earth (Sánchez-Baracaldo, et al. 2017). According to recent classifications there are five divisions of archaeplastida, the rhodophytes (red algae), glaucophytes, Rhodelphidia, Picozoa and Viridiplantae (green plants) (Bowles, et al. 2022).

Red algae are comprised of about 7,000 morphologically quite diverse species including mostly multicellular specimens that occur in aquatic habitats (Yoon, et al. 2010; Dittami, et al. 2017). Rhodelphidia and Picozoa are recently identified phyla including only a few species which are non-photosynthetic and live in aquatic environments (Gawryluk, et al. 2019; Schön, et al. 2021). Glaucophytes are also a small group comprised of about 20 species of unicellular algae that are found in freshwater environments (Figueroa-Martinez, et al. 2019).

The largest division of Archaeplastida are the Viridiplantae that arose about 1 billion years ago and are characterized by complex cell walls (Leliaert, et al. 2011). There are approximately 500,000 species of green plants (One Thousand Plant Transcriptomes Initiative 2019) that are found in aquatic as well as in terrestrial environments (Leliaert, et al. 2011). Green plants are further subdivided into chlorophytes, prasinodermophytes and streptophytes (Leliaert, et al. 2011). Chlorophytes are mainly unicellular but display a large diversity of body forms. There are about 4,300 species of chlorophytes which inhabit freshwater environments (Guiry 2012). Prasinodermophytes are a group of aquatic organisms that have been recently suggested as a new phylum in green plants (Li, et al. 2020). Within the green plants, the streptophytes are the largest group (Leliaert, et al. 2011). The ancestor of streptophytes was most likely multicellular (Bowles, et al. 2022) from which unicellular and multicellular species evolved. Streptophytes are comprised of five paraphyletic lineages of streptophyte algae and embryophytes (land plants) (Leliaert, et al. 2011). The streptophyte algae classes Chlorokybophyceae and Mesostigmatophycae form a clade and branched off first, followed by the Klebsormidiophyceae, Charophyceae and the Coleochaetophyceae (Bowles, et al. 2022). Streptophyte algae of the class Zygnematophyceae are likely the closest relatives of land plants.

### Evolution of MADS-box genes

In land plants, the evolution of MADS-box genes has been well documented. While the genomes of early-branching land plants like mosses or spike-mosses contain around 20 MADS-box genes, this number increases to more than 100 genes in flowering plant genomes (Gramzow, et al. 2010; One Thousand Plant Transcriptomes Initiative 2019). Thereby, both types of MADS-box genes have expanded. However, it has been noticed that Type I and Type II genes in plants have dramatically different evolutionary dynamics (Gramzow and Theißen 2013). For example, Type I genes experience faster birth and death rates than Type II genes (Nam, et al. 2004); Type II genes are much more conserved but steadily increased in number mainly by the preferential retention and diversification of gene copies after whole genome duplications (Theißen and Gramzow 2016). The increase in the number of especially Type II MADS-box genes is correlated with and probably even causally linked to the increasing complexity of the body plans of land plants (Theissen, et al. 2000; Gramzow and Theissen 2010).

Much less is known about MADS-box genes in rhodophyte, chlorophyte and streptophyte algae. The genomes of the red algae species *Porphyra umbilicalis* and *P. purpurea* each have been found to contain two MADS-box genes (Stiller, et al. 2012). While one of the genes is expressed about equally in gametophytic and sporophytic life stages, the other one shows lower expression in the gametophyte than in the sporophyte. Only one MADS-box gene has been identified in the red alga *Cyanidioschyzon merolae* (Stiller, et al. 2012) and in *Galdiera* sp. (Hirooka, et al. 2022). In *Galdiera* sp. the MADS-box gene is specifically expressed in the sporophyte and has been hypothesized to be involved in the transition from gametophyte to sporophyte. All of these MADS-box genes from red algae are proposed to be Type II but no K box was reported (Stiller, et al. 2012; Hirooka, et al. 2022).

One MADS-box gene of a chlorophyte has been described in some detail: *CsubMADS1* from *Coccomyxa subellipsoidea* (Nayar and Thangavel 2021). This gene is expressed in the lag phase of growth and has been hypothesized to confer stress tolerance during this phase. Like the described MADS-box genes of red algae, *CsubMADS1* has also been suggested to be a Type II gene that does not, however, encode a K domain (Nayar and Thangavel 2021).

MADS-box genes have also been identified relatively early in the streptophyte algae *Chara globularis, Coleochaete scutata*, and *Closterium peracerosum–strigosum–littorale* complex (Tanabe, et al. 2005). Expression of these genes has been detected mainly in the gametophyte during differentiation into reproductive structures and hence, it has been hypothesized that they function in haploid reproductive cell differentiation (Tanabe, et al. 2005). The MADS-box genes of the streptophyte algae have also been assigned to the Type II genes but unlike the aforementioned genes of rhodophyte and chlorophyte algae, they also encode a K domain (Tanabe, et al. 2005). A more recent whole genome analysis of *Chara braunii* identified three Type II genes; while one does not encode a K domain, the two others are of the MIKC-type, even though one gene has a non-canonical exon-intron structure (Nishiyama, et al. 2018).

Within Archaeplastida MIKC-type genes hence likely evolved by the acquisition of the K box in the stem group of streptophytes, that is after chlorophytes but before all of the streptophyte algae lineages branched off. Most authors assume that the K box was gained by a Type II gene (Alvarez-Buylla, et al. 2000; Kaufmann, et al. 2005; Tanabe, et al. 2005). However, there is also one study that claims that the K box was acquired by a Type I gene (Lai, et al. 2020).

Hence, how the way was paved for Type II MADS-box genes to gain their major functions in the morphological development in land plants during the early evolution of Archaeplastida remains largely unknown. Here, we make use of the increasing amount of transcriptomic and genomic data for early-branching Archaeplastida to study the early evolution of MADS-box genes in this eukaryotic supergroup. We find that archaeplastid algae likely only feature Type II genes which remained largely single copy until streptophytes emerged. In the stem group of streptophytes, Type II genes were duplicated and one of the duplicates acquired a K box, giving rise to MIKC-type genes. The clade not possessing the K box was lost in the stem group of land plants after streptophyte algae branched-off, leaving MIKC-type genes to evolve into almost ubiquitous regulators of plant development.

## Results

We have identified one to five MADS-box genes in all investigated species of rhodophyte, glaucophyte, chlorophyte, prasinodermophyte and streptophyte algae (Supplementary table 1). For most species, we found one MADS-box gene and only rarely more than two MADS-box genes were identified from one species.

Next, we checked the domain structure of the encoded algae proteins using the conserved domains database (CDD) search tool on NCBI (Lu, et al. 2020). For 27 proteins, the search revealed a coupling of the MADS with a K domain (Supplementary table 2). One protein additionally contained a G-protein-pathway-suppressor domain. The K domain was identified in MADS-domain proteins of all classes of streptophyte algae except Mesostigmatophyceae, but not in any other algae. This suggests that a MADS-box gene was recombined with a K box in the stem group of streptophytes.

To elucidate to which type the newly identified MADS-box genes from rhodophyte, glaucophyte, chlorophyte, prasinodermophyte and streptophyte algae belong, we reconstructed their phylogeny. To obtain a better resolution, we included MADS-box genes from land plants and from eukaryotic supergroups other than the Archaeplastida (Supplementary table 1). Apart from two MADS-box genes from the chlorophyte *Aphanochaete repens*, we did not find any Type I MADS-box gene in the algae investigated by us (Supplementary figure 1). These two genes cluster within a clade of SRF-like MADS-box genes from Amorphea, Cryptista and Haptista, but are not closely related to any plant MADS-box gene. They may represent contamination or horizontally transferred genes and hence we ignored them in our further analyses.

The lack of Type I MADS-box genes in the algae studied here is supported by Multi-Harmony analysis. For this analysis, we took our alignment of the MADS domain and removed all sequences that were not from fungi or animals. We defined the groups of Type I and Type II MADS domains based on public annotations, if available, or on phylogenetic relatedness to annotated domains in our phylogeny. The Multi-Harmony analyses identified 18 positions in our alignment that distinguish Type I and Type II MADS domains from animals and fungi. Investigating the orthologous positions in MADS domains of archaeplastid algae, only two sequences contained more residues corresponding to the consensus of Type I than to the consensus of Type II (Supplementary table 3). These two sequences were the ones from the chlorophyte *Aphanochaete repens* which also clustered with Type I genes in our phylogeny.

We reconstructed another phylogeny only including Type II genes from Archaeplastida as identified in our larger phylogeny. Our phylogeny suggests that there was likely a single (Type II) MADS-box gene in the stem group of Archaeplastida (Figure 1). The number of MADS-box genes remained low in rhodophytes, glaucophytes, prasidermatophytes as well as in chlorophytes with few, mainly recent duplications. In streptophyte algae, in contrast, there are at least two groups of MADS-box genes (Figure 1). Interestingly, one of these groups contains Type II genes not encoding a K domain while the other group consist of genes containing a K box.

**Figure 1:**
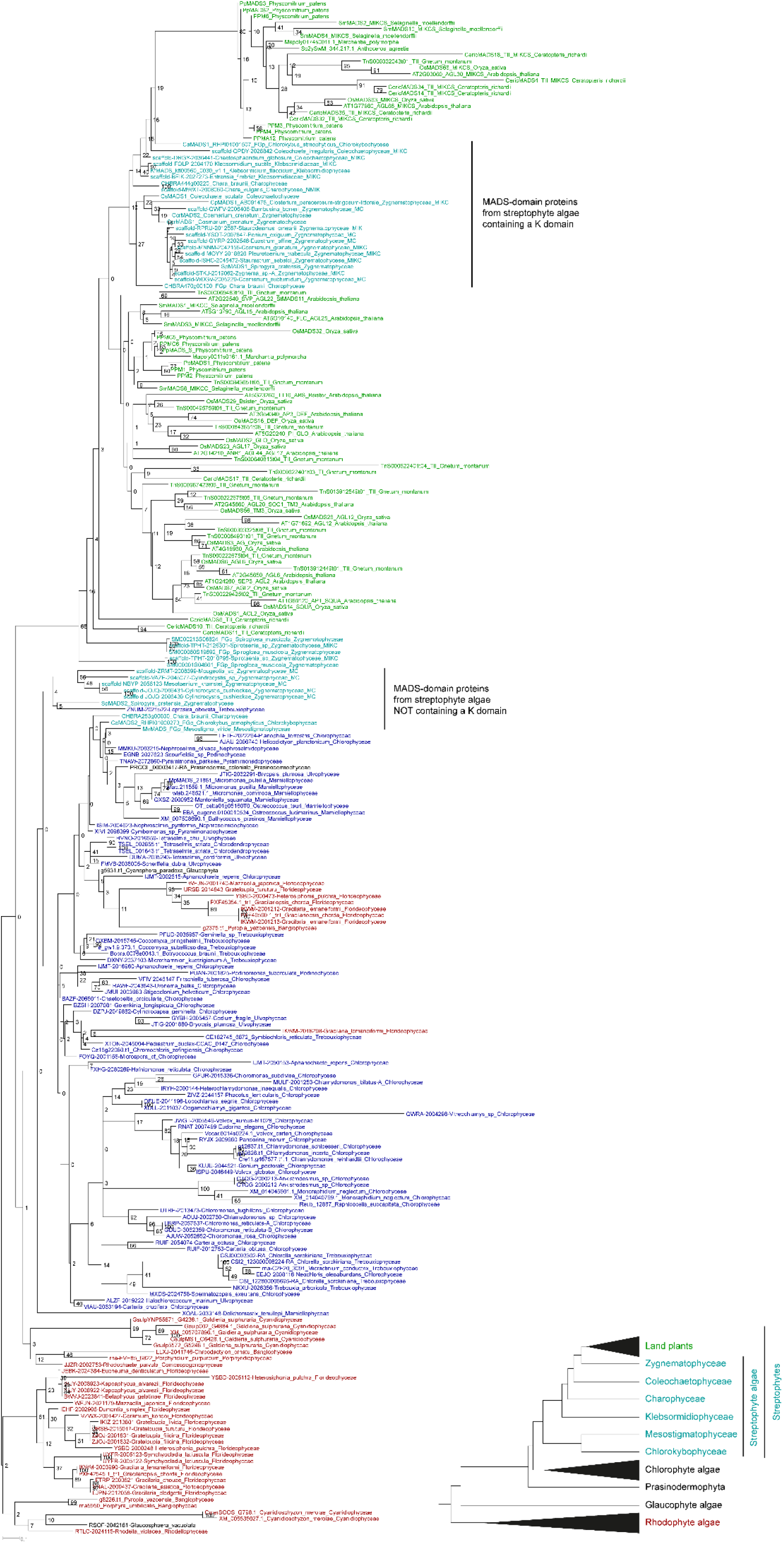
Phylogeny of Type II MADS-domain proteins from Archaeplastida. The phylogeny was reconstructed using RAxML based on an alignment of the MADS domains and the first approximately 15 amino acids of the I domain. Names of MADS-domain proteins are colored according to the species in which they were identified as indicated in the species phylogeny on the bottom right. The two groups of MADS-domain proteins from streptophyte algae, one with and one without the K domain, are indicated in the phylogeny. Numbers on the nodes reflect bootstrap values of 1000 bootstrap replicates.

To reveal sequence differences between the MADS domains of these two groups, we conducted another Multi-Harmony analysis. We aligned complete sequences of MADS-domain proteins from streptophyte algae and removed all positions that are not part of the MADS domain. We defined two groups based on the presence or absence of a K domain in the remainder of the protein. The Multi-Harmony analysis identified 17 positions that distinguish the MADS domains of these two groups (Figure 2, Supplementary table 4), suggesting that MADS-box genes from streptophyte algae indeed constitute two well separable groups. This suggests that there has been a duplication of MADS-box genes in the stem group of extant streptophytes (Figure 3). One of the duplicates acquired a K box giving rise to MIKC genes. Both versions, with and without a K box, were retained in all classes of extant streptophyte algae except Mesostigmatophyceae. In land plants, no Type II genes without a K box are known, indicating that the Type II genes without a K box were lost in the stem group of land plants (Figure 3).

**Figure 2:**
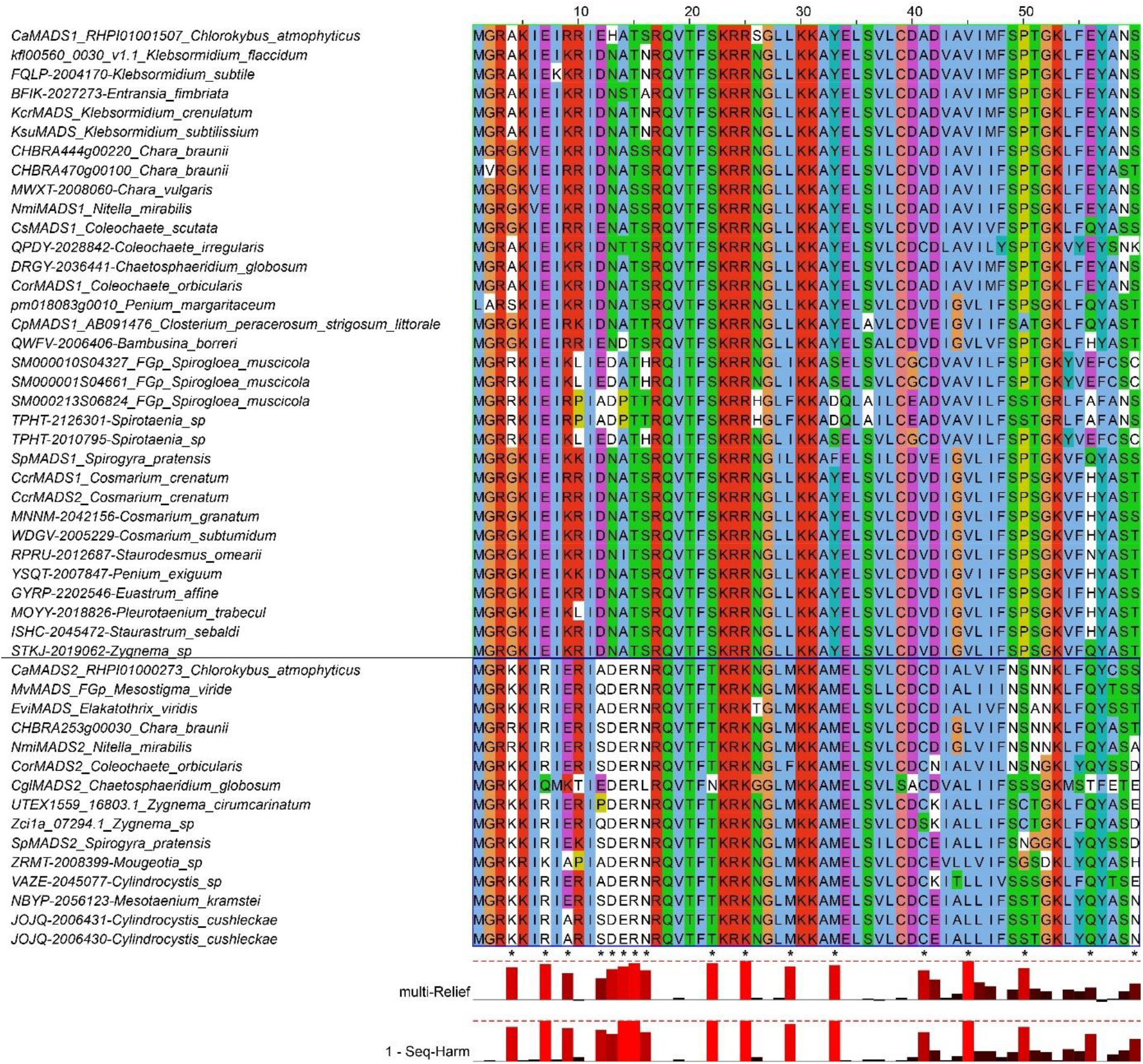
Alignment of MADS domains from streptophyte algae. Complete protein sequences were aligned using MAFFT with the L-INS-I option that is optimized to align a set of sequences containing one alignable domain surrounded by unalignable sequences. The alignment was manually reduced to the MADS domain and groups were defined based on whether the complete proteins include the K domain (upper group) or not (lower group). The alignment was then subjected to MultiHarmony analysis that identified 17 positions (marked by “*”) in the alignment that distinguish these two groups.

**Figure 3:**
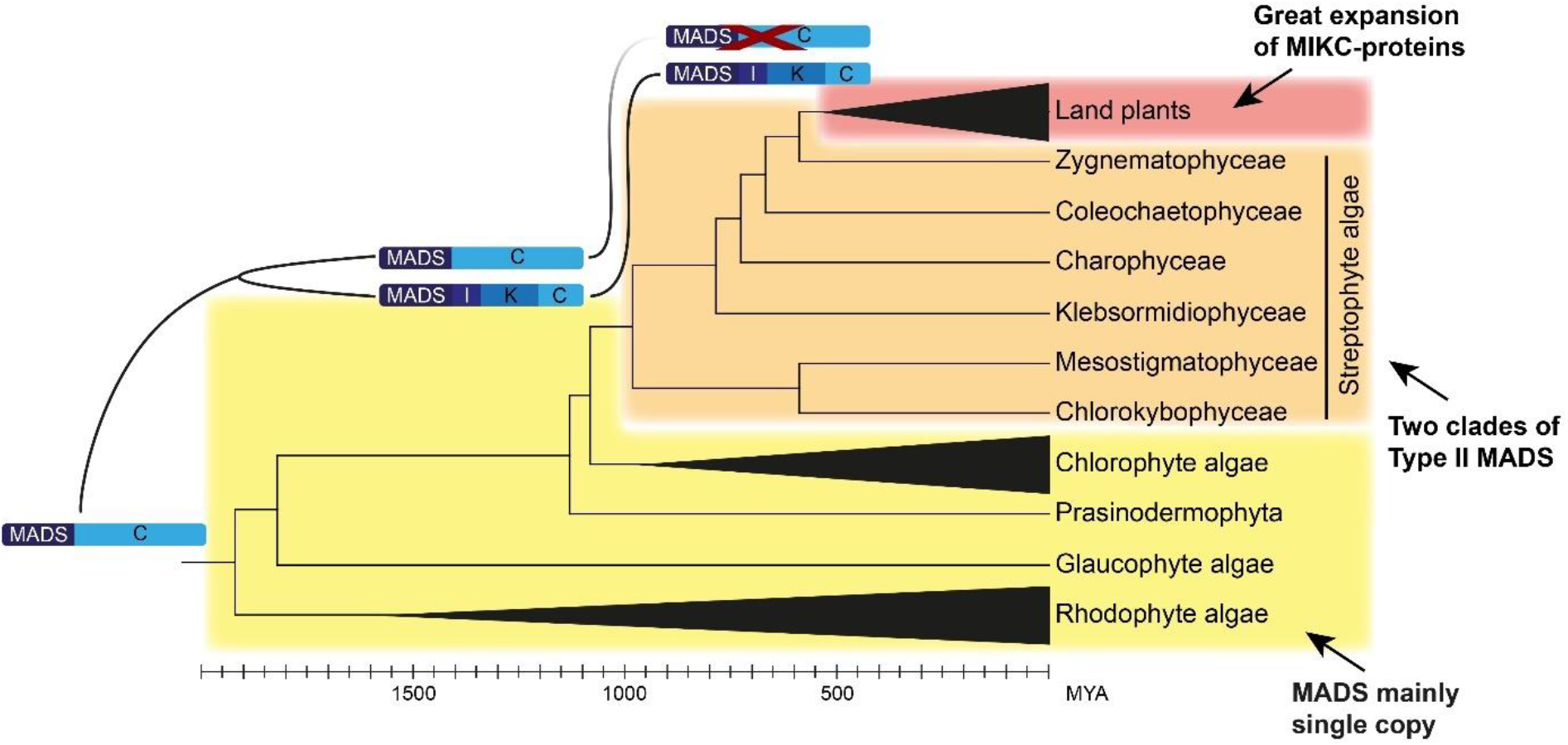
Scenario for the early evolution of MADS-box genes in Archaeplastida. Events and patterns in the evolution of MADS-box genes are mapped to the phylogeny of Archaeplastida. The ancestor of Archaeplastida likely possesed one Type II MADS-box gene without a K box. MADS-box genes consequently remained mainly single copy in rhodophyte, glaucophyte, prasinodermophyte and chlorophyte algae and their ancestors (yellow shading). In an ancestor of streptophytes, after chlorophytes branched off, the MADS-box gene was duplicated and one of the duplicates aquired the K box. In streptophyte algae and their ancestor there were hence two clades of Type II genes (orange shading). At the base of land plants, the clade of MADS-box genes without a K box was lost and the other clade of MIKC genes greatly expanded (red shading).

## Discussion

Our study represents the first comprehensive inventory of MADS-box genes of archaeplastid algae. Apart from two possible contaminant or horizontally transferred genes, no Type I gene has been identified in archaeplastid algae (Figure 1, Supplementary table 3). This is in line with a recent preprint that suggests that the Type I MADS-box genes of land plants evolved by a duplication of an Type II gene within land plants (Qiu, et al. 2023).

The MADS-box genes we identified in rhodophyte, glaucophyte, prasinodermophyte and chlorophyte algae do not encode a K domain (Supplementary table 2). Nevertheless, our phylogeny analyses (Figure 1) indicate that these genes belong to the clade of Type II genes. This is in contrast to previous publications that interpreted them as Type I genes (Gramzow and Theissen 2010; Barker and Ashton 2013) but in line with a more recent publication (Nayar and Thangavel 2021). MADS-box genes containing a K box were solely identified from streptophytes, where already the genome of one of the most basal classes of streptophyte algae, Chlorokybophyceae, contains a MIKC gene. This corroborates earlier hypotheses (Gramzow and Theissen 2010; Nayar and Thangavel 2021) and reveals that the K box was gained by a Type II MADS-box gene in an ancestor of streptophytes after chlorophytes branched off. MIKC genes, more specifically the subclade of MIKC^C^ genes, are well known for their crucial function in the development especially of flowering plants (Gramzow and Theissen 2010; Smaczniak, et al. 2012). An important aspect that may have allowed these genes to take over functions in the determination of different kinds of structures, such as the floral organs, is likely the ability of the encoded proteins to form tetrameric complexes such as the floral quartets (Theissen and Saedler 2001; Theißen, et al. 2016). Tetramerization of MIKC^C^ proteins is conferred by the C-terminal part of the K domain (Puranik, et al. 2014). Recently, however, it has been shown that MIKC proteins from some streptophyte algae do only have a weak or no ability to form tetrameric complexes, this way corroborating the view that the K domain may initially have been mainly involved in strengthening protein dimerization (Kaufmann, et al. 2005; Rümpler, et al. 2022). Nevertheless, the acquisition of the K box by a MADS-box gene in an ancestor of streptophytes may represent some kind of pre-adaptation that, after an elongation of the part encoding the C-terminal region of the K domain (Rümpler, et al. 2022), gained the ability of tetramerization. Tetramerization may then have enabled the specific recognition of diverse target genes. Sequence diversification of paralogous copies of MIKC-type genes may subsequently have enabled land plants to develop specialized structures that boosted complexity, disparity and adaptability of plants on land, eventually leading to well-known evolutionary novelties such as seeds, flowers and fruits.

We have identified at least one MADS-box gene in all taxonomic classes of archaeplastid algae (Supplementary table 1), suggesting an essential function for MADS-box genes in these algae. Deletion of the only MADS-box gene of *Galdiera* sp. prevents these red algae from progressing from gametophyte to sporophyte (Hirooka, et al. 2022) while overexpression of the MADS-box gene of the chlorophyte green algae *Coccomyxa subellipsoidea* leads to formation of polyploid multinucleate cells (Nayar and Thangavel 2021). Based on this very limited functional data, one may hypothesize that MADS-domain proteins in archaeplastid algae are involved in basic cell-biological processes like cell-cycle progression as has been observed for some MADS-box genes in animals and fungi (Elble and Tye 1992; Badodi, et al. 2015) or basic cell-differentiation processes as in amoeba and animals (Galardi-Castilla, et al. 2013; Wang, et al. 2018).

The number of identified MADS-box genes per species as well as our phylogeny reconstruction suggests that these genes remained mainly single copy in the ancestor and during the evolution of rhodophyte, glaucophyte, prasinodermophyte and chlorophyte algae (Figure 3). At the base of streptophytes, however, the MADS-box gene was likely duplicated and one of the duplicates acquired a K box as second conserved sequence element. This resulted in two groups of Type II MADS-box genes in streptophyte algae, one with and one without a K box (Figure 2). Hence, there was a slight increase in the number of MADS-box genes at the base of streptophytes. However, this increase was not intensified in streptophyte algae by further duplications. Most remarkably, the group not encoding the K domain was lost at the base of land plants (Figure 3). Also in the stem group of land plants, MIKC-type genes were duplicated to give rise to the two clades of MIKC^C^ and MIKC* genes (Rümpler, et al. 2022), followed by a great expansion mainly of the MIKC^C^ genes and the evolution of a diversity of functions that the encoded proteins are so well known for in flowering plants, ranging from root to flower and fruit development (Gramzow and Theissen 2010; Smaczniak, et al. 2012).

Our study shows that Type II MADS-box genes of rhodophytes, glaucophytes, prasinodermophytes and chlorophytes do not encode a K domain - hence they are not of the MIKC-type - and remained mainly single-copy in these lineages. The ancestral Type II MADS-box gene was duplicated and the K box was recombined with one of the duplicates in an ancestor of streptophytes after chlorophytes branched off. The transition to land of plants was then accompanied by a loss of the ancestral, non-MIKC Type II gene not encoding a K domain and duplication of the MIKC-type gene, yielding MIKC^C^ and MIKC*. Hence, our study provides important new insights into the deep evolution of Type II MADS-box genes in Archaeplastida that paved the way for MADS-domain proteins to become a crucial family of transcription factors in land plants, and for land plants to become the arguably most crucial general component of terrestrial ecosystems.

## Materials and methods

### Data collection

Detailed information on the databases from which MADS-domain proteins of different species were retrieved, are given in Supplementary table 1.

Shortly, first MADS-domain proteins were compiled from published datasets (Gramzow, et al. 2012; One Thousand Plant Transcriptomes Initiative 2019; Marchant, et al. 2022). Additional MADS-box genes from streptophyte algae were identified by BLAST searches of transcriptome datasets available at the Sequence Read Archive (SRA) at the National Centre for Biotechnology information (NCBI) (Sayers, et al. 2023). MADS-box sequences of other streptophyte algae were used as search sequences and the identified reads were assembled using Sequencher (Sequencher 2020). The ends of the resulting contig were iteratively used as search sequences to extend the contig until the coding sequence was completely assembled or no more sequences were found.

MADS-domain proteins from other species were identified by searching for proteins annotated to contain the PFAM domain “PF00319” in PhycoCosm (Grigoriev, et al. 2021), MycoCosm (Ahrendt, et al. 2023) and Ensembl (Yates, et al. 2022).

### Phylogeny reconstruction

MADS-domain proteins were aligned using MAFFT with the “`L-INS-I`” option (Katoh, et al. 2019). The alignment was manually inspected to exclude proteins with poor fit to the MADS domain from the dataset. The reduced dataset was subsequently aligned again using MAFFT with the same option. The novel alignment was manually trimmed to only leave positions corresponding to the MADS domain and the following about 15 amino acids which may represent a common intervening (I) domain (Lai, et al. 2021). Based on the trimmed alignment, the phylogeny was reconstructed using RAxML (Stamatakis 2014) on the CIPRES Science Gateway (Miller, et al. 2011) choosing the LG amino acid substitution matrix (Le and Gascuel 2008) and generating 1000 rapid bootstrap replicates. Based on the complete phylogeny, Type II MADS-domain proteins from Archaeplastida were determined. The Type II MADS-domain proteins from Archaeplastida were aligned and their phylogeny was reconstructed as described for the whole dataset. The protein phylogenies inferred this way were taken as a proxy for the evolution of the encoding genes.

### Multi-Harmony analyses

First, we aimed to identify positions in the MADS domain that distinguish Type I and Type II MADS domains, where we concentrated on proteins from animals and fungi. Therefore, the trimmed alignment as build for the phylogeny reconstruction of the complete dataset was taken and all sequences not from animals and fungi were removed. Groups of Type I and Type II MADS domains were then defined based on previous annotations or relatedness to previously annotated domains in our phylogeny. Multi-Harmony analysis (Brandt, et al. 2010) was started in Jalview (Procter, et al. 2021) and the results were obtained from the website http://zeus.few.vu.nl. As used by the authors of the Multi-Harmony method, we considered positions with a sequence-harmony score of less than 0.5 and a multi-relief weight of more than 0.8 to distinguish between the two groups.

To distinguish MADS domains from proteins of streptophyte algae with and without a K domain, another Multi-Harmony analysis was conducted. The sequences of MADS-domain proteins of streptophyte algae were aligned using MAFFT as described above. The alignment was manually trimmed to only contain the MADS domain. Two groups were defined based on whether the complete protein sequences contained a K domain or not. The actual Multi-Harmony analysis (Brandt, et al. 2010) was conducted as explained above.

## Acknowledgements

We thank all international sequencing and annotation efforts giving us plenty of data to analyse.

