## Supplementary figures and images for "Deep evolution of MADS-box genes in Archaeplastida"

### Supplementary figure 1

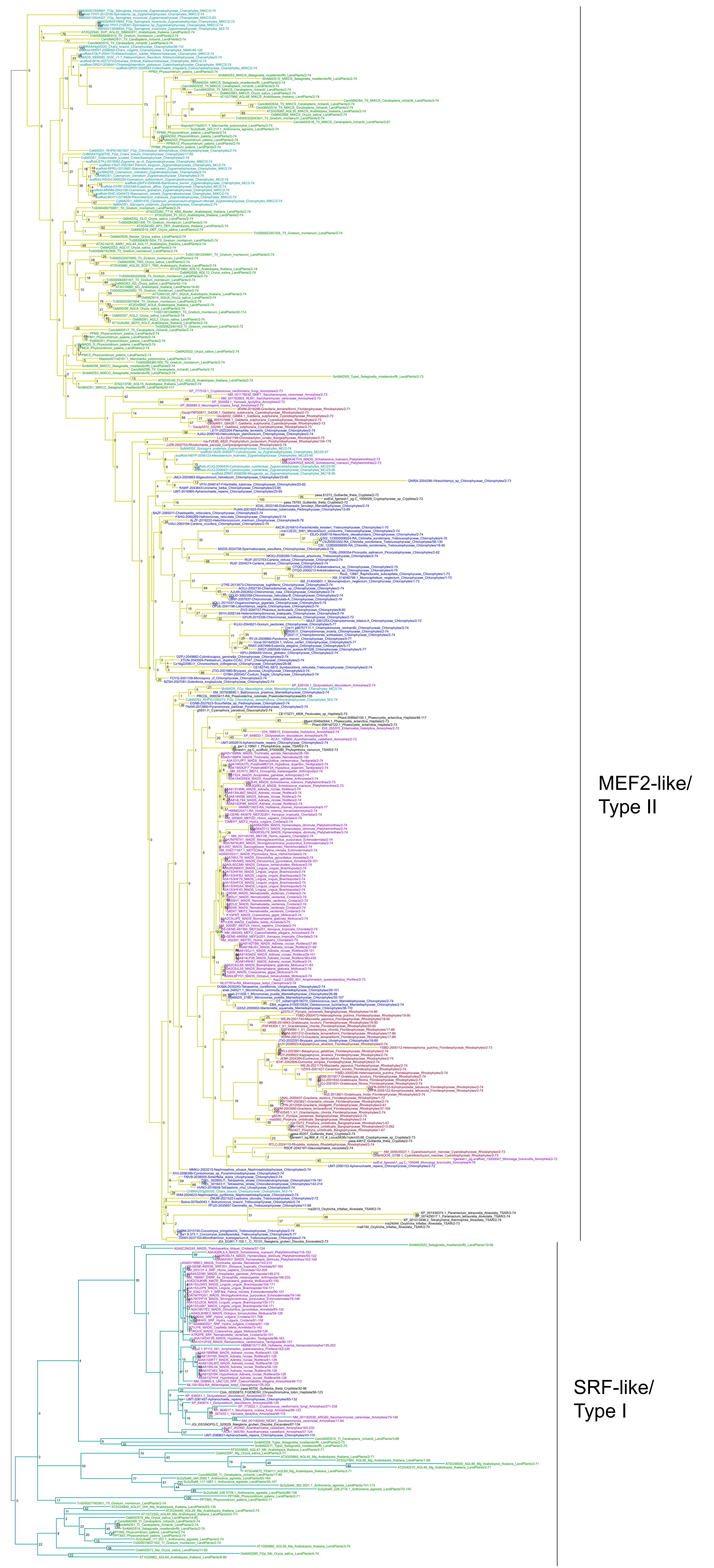
