## Supplementary table 1 for "Deep evolution of MADS-box genes in Archaeplastida"

### Data access

| Supergroup | Division | Species | Data source | Total MADS |
|---|---|---|---|---|
| Archaeplastida | Red algae | *Galdieria sulphuraria* | PhycoCosm | 5 |
| | | *Cyanidioschyzon merolae* | PhycoCosm | 2 |
| | | *Rhodochaete parvula* | 1KP | 1 |
| | | *Porphyridium purpureum* | PhycoCosm | 1 |
| | | *Rhodella violacea* | 1KP | 1 |
| | | *Betaphycus gelatinae* | 1KP | 1 |
| | | *Gracilaria chouae* | 1KP | 1 |
| | | *Gracilaria lemaneiformi* | 1KP | 4 |
| | | *Gracilaria blodgettii* | 1KP | 1 |
| | | *Gracilaria asiatica* | 1KP | 1 |
| | | *Dumontia simplex* | 1KP | 1 |
| | | *Kappaphycus alvarezii* | 1KP | 2 |
| | | *Grateloupia livida* | 1KP | 1 |
| | | *Grateloupia turuturu* | 1KP | 2 |
| | | *Grateloupia filicina* | 1KP | 2 |
| | | *Eucheuma denticulatum* | 1KP | 1 |
| | | *Symphyocladia latiuscula* | 1KP | 2 |
| | | *Ceramium kondoi* | 1KP | 1 |
| | | *Mazzaella japonica* | 1KP | 2 |
| | | *Heterosiphonia pulchra* | 1KP | 3 |
| | | *Gracilariopsis chorda* | PhycoCosm | 3 |
| | | *Glaucosphaera vacuolata* | 1KP | 1 |
| | Glaucophytes | *Cyanophora paradoxa* | PhycoCosm | 1 |
| | Green plants (chlorophytes) | *Nephroselmis pyriformis* | 1KP | 1 |
| | | *Nephroselmis olivace* | 1KP | 1 |
| | | *Pyramimonas parkeae* | 1KP | 1 |
| | | *Cymbomonas sp* | 1KP | 1 |
| | | *Micromonas pusilla* | PhycoCosm | 2 |
| | | *Micromonas commoda* | PhycoCosm | 1 |
| | | *Mantoniella squamata* | 1KP | 1 |
| | | *Dolichomastix tenuilepi* | 1KP | 1 |
| | | *Bathycoccus prasinos* | PhycoCosm | 1 |
| | | *Ostreococcus lucimarinus* | PhycoCosm | 1 |

| | | | | |
|---|---|---|---|---|
| | | *Ostreococcus tauri* | PhycoCosm | 1 |
| | | *Picocystis salinarum* | 1KP | 1 |
| | | *Scourfieldia sp* | 1KP | 1 |
| | | *Pedinomonas tuberculata* | 1KP | 1 |
| | | *Tetraselmis striata* | PhycoCosm | 2 |
| | | *Tetraselmis cordiformis* | 1KP | 1 |
| | | *Tetraselmis chui* | 1KP | 1 |
| | | *Botryococcus braunii* | PhycoCosm | 1 |
| | | *Parachlorella kessleri* | 1KP | 1 |
| | | *Microthamnion kuetzigianum* | 1KP | 1 |
| | | *Coccomyxa pringsheimii* | 1KP | 1 |
| | | *Coccomyxa subellipsoidea* | PhycoCosm | 1 |
| | | *Trebouxia arboricola* | 1KP | 1 |
| | | *Geminella sp* | 1KP | 1 |
| | | *Leptosira obovata* | 1KP | 1 |
| | | *Chlorella sorokiniana* | PhycoCosm | 3 |
| | | *Micractinium conductrix* | PhycoCosm | 1 |
| | | *Symbiochloris reticulata* | PhycoCosm | 1 |
| | | *Monoraphidium neglectum* | PhycoCosm | 2 |
| | | *Raphidocelis subcapitata* | PhycoCosm | 1 |
| | | *Chromochloris zofingiensis* | PhycoCosm | 1 |
| | | *Ankistrodesmus sp* | 1KP | 2 |
| | | *Aphanochaete repens* | 1KP | 5 |
| | | *Carteria crucifera* | 1KP | 1 |
| | | *Carteria obtusa* | 1KP | 2 |
| | | *Chaetopeltis orbicularis* | 1KP | 1 |
| | | *Chlamydomonas reinhardtii* | PhycoCosm | 1 |
| | | *Chlamydomonas incerta* | PhycoCosm | 1 |
| | | *Chlamydomonas schloesseri* | PhycoCosm | 1 |
| | | *Chlamydomonas bilatus* | 1KP | 1 |
| | | *Chlamydomonas sp* | 1KP | 1 |
| | | *Chloromonas reticulata* | 1KP | 1 |
| | | *Chloromonas rosa* | 1KP | 1 |
| | | *Chloromonas subdivisa* | 1KP | 1 |
| | | *Chloromonas tughillensi* | 1KP | 1 |
| | | *Cylindrocapsa geminella* | 1KP | 1 |
| | | *Eudorina elegans* | 1KP | 1 |

| | | | | |
|---|---|---|---|---|
| | | *Fritschiella tuberosa* | 1KP | 1 |
| | | *Golenkinia longispicula* | 1KP | 1 |
| | | *Gonium pectorale* | 1KP | 1 |
| | | *Hafniomonas reticulata* | 1KP | 1 |
| | | *Helicodictyon planctonicum* | 1KP | 1 |
| | | *Heterochlamydomonas inaequalis* | 1KP | 1 |
| | | *Lobochlamys segnis* | 1KP | 1 |
| | | *Microspora cf* | 1KP | 1 |
| | | *Neochloris oleoabundans* | 1KP | 1 |
| | | *Oogamochlamys gigantea* | 1KP | 1 |
| | | *Pandorina morum* | 1KP | 1 |
| | | *Pediastrum duplex* | 1KP | 1 |
| | | *Phacotus lenticularis* | 1KP | 1 |
| | | *Planophila terrestris* | 1KP | 1 |
| | | *Spermatozopsis exsultans* | 1KP | 1 |
| | | *Stigeoclonium helveticum* | 1KP | 1 |
| | | *Uronema belka* | 1KP | 1 |
| | | *Vitreochlamys sp* | 1KP | 1 |
| | | *Volvox aureus* | 1KP | 1 |
| | | *Volvox globator* | 1KP | 1 |
| | | *Volvox carteri* | PhycoCosm | 1 |
| | | *Halochlorococcum marinum* | 1KP | 1 |
| | | *Scherffelia dubia* | 1KP | 1 |
| | | *Codium fragile* | 1KP | 1 |
| | | *Bryopsis plumosa* | 1KP | 2 |
| | Green plants (Prasinodermatophyta) | *Prasinoderma coloniale* | PhycoCosm | 1 |
| | Green plants (streptophytes) | *Chlorokybus atmophyticus* | PhycoCosm | 2 |
| | | *Mesostigma viride* | PhycoCosm | 1 |
| | | *Klebsormidium nitens* | PhycoCosm | 1 |
| | | *Klebsormidium subtile* | 1KP | 1 |
| | | *Entransia fimbriata* | 1KP | 1 |
| | | *Chara braunii* | PhycoCosm | 3 |
| | | *Chara vulgaris* | 1KP | 1 |
| | | *Coleochaete scutata* | NCBI SRA | 1 |
| | | *Coleochaete irregularis* | 1KP | 1 |

| | | | | |
|---|---|---|---|---|
| | | *Chaetosphaeridium globosum* | 1KP | 1 |
| | | *Spirogloea muscicola* | PhycoCosm | 4 |
| | | *Spirogyra pratensis* | NCBI SRA | 2 |
| | | *Cosmarium crenatum* | NCBI SRA | 2 |
| | | *Cosmarium subtumidum* | 1KP | 1 |
| | | *Cosmarium granatum* | 1KP | 1 |
| | | *Closterium peracerosum-strigosum-littorale* | NCBI SRA | 1 |
| | | *Bambusina borreri* | 1KP | 1 |
| | | *Euastrum affine* | 1KP | 1 |
| | | *Pleurotaenium trabecula* | 1KP | 1 |
| | | *Staurastrum sebaldi* | 1KP | 1 |
| | | *Staurodesmus omearii* | 1KP | 1 |
| | | *Penium exiguum* | 1KP | 1 |
| | | *Zygnema sp.* | 1KP | 1 |
| | | *Spirotaenia sp* | 1KP | 2 |
| | | *Cylindrocystis cushleckae* | 1KP | 2 |
| | | *Cylindrocystis sp* | 1KP | 1 |
| | | *Mesotaenium kramstei* | 1KP | 1 |
| | | *Mougeotia sp* | 1KP | 1 |
| | | *Marchantia polymorpha* | Phytozome | |
| | | *Physcomitrium patens* | Phytozome | |
| | | *Selaginella moellendorffii* | Phytozome | |
| | | *Anthoceros agrestis* | NCBI Genome | |
| | | *Ceratopteris richardii* | Phytozome | |
| | | *Gnetum montanum* | NCBI Genome | |
| | | *Oryza sativa* | Phytozome | |
| | | *Arabidopsis thaliana* | TAIR | |
| Cryptista | | *Guillardia theta* | PhycoCosm | 5 |
| | | *Cryptophyceae sp* | PhycoCosm | 2 |
| Haptista | | *Chrysochromulina tobin* | PhycoCosm | 1 |
| | | *Phaeocystis antarctica* | PhycoCosm | 3 |
| | | *Pavlovales sp* | PhycoCosm | 1 |
| TSAR | | *Phytophthora sojae* | MycoCosm | 1 |
| | | *Phytophthora ramorum* | PhycoCosm | 1 |
| | Alveolata | *Tetrahymena thermophila* | NCBI Genbank | 1 |
| | | *Paramecium tetraurelia* | NCBI Genbank | 2 |

| | | | | |
|---|---|---|---|---|
| | | *Oxytricha trifallax* | MycoCosm | 3 |
| Amorphea | Amoebozoa | *Entamoeba histolytica* | Ensembl | 2 |
| | | *Acanthamoeba castellanii* | Ensembl | 3 |
| | | *Dictyostelium discoideum* | NCBI Genbank | 4 |
| | Fungi | *Saccharomyces cerevisiae* | NCBI Genbank | 4 |
| | | *Cryptococcus neoformans* | MycoCosm | 2 |
| | | *Yarrowia lipolytica* | NCBI Genbank | 2 |
| | | *Neurospora crassa* | NCBI Genbank | 2 |
| | Animals (Cnidaria) | *Thelohanellus kitauei* | Ensembl | 1 |
| | | *Nematostella vectensis* | Ensembl | 7 |
| | | *Hydra vulgaris* | Ensembl | 4 |
| | Animals (Ctenophora) | *Mnemiopsis leidyi* | Ensembl | 2 |
| | Animals (Porifera) | *Amphimedon queenslandica* | Ensembl | 2 |
| | Animals (Bilateria) | *Strongylocentrotus purpuratus* | Ensembl | 4 |
| | | *Patiria miniata* | Ensembl | 2 |
| | | *Saccoglossus kowalevskii* | Ensembl | 1 |
| | | *Ptychodera flava* | UniProt | 1 |
| | | *Biomphalaria glabrata* | Ensembl | 4 |
| | | *Crassostrea gigas* | Ensembl | 3 |
| | | *Octopus bimaculoides* | Ensembl | 3 |
| | | *Trichinella spiralis* | Ensembl | 3 |
| | | *Caenorhabditis elegans* | NCBI Genbank | 2 |
| | | *Hymenolepis diminuta* | Ensembl | 5 |
| | | *Schistosoma mansoni* | Ensembl | 5 |
| | | *Adineta ricciae* | Ensembl | 19 |
| | | *Hypsibius dujardini* | Ensembl | 3 |
| | | *Ramazzottius varieornatus* | UniProt | 2 |
| | | *Hofstenia miamia* | Ensembl | 3 |
| | | *Capitella teleta* | Ensembl | 2 |
| | | *Dimorphilus gyrociliatus* | Ensembl | 3 |
| | | *Lingula unguis* | Ensembl | 11 |
| | | *Drosophila melanogaster* | NCBI Genbank | 2 |
| | | *Anopheles gambiae* | Ensembl | 3 |
| | | *Homo sapiens* | NCBI Genbank | 5 |
| | | *Xenopus tropicalis* | Ensembl | 4 |
| | Choanozoa | *Monosiga brevicollis* | MycoCosm | 2 |
| Excavates | | *Naegleria gruberi* | PhycoCosm | 2 |
